# Electrophysiological signature of the interplay between habits and inhibition in response to smoking-related cues in individuals with a smoking habit: an ERP study

**DOI:** 10.1101/2022.07.28.501841

**Authors:** Julien Dampuré, Paola Agudelo-Orjuela, Maartje Van Der Meij, David Belin, Horacio A. Barber

**Author notes:** corresponding author Address for correspondence: Julien Dampuré, PhD, Facultad de Psicología, Universidad de La Sabana, Chía, Colombia. co-last authors (David Belin, Horacio A. Barber).

## Abstract

The rigid, stimulus-bound nature of drug seeking that characterizes Substance-use disorder (SUD) has been related to a dysregulation of motivational and early attentional reflexive and inhibitory reflective systems. However, the mechanisms by which these systems are engaged by drug-paired conditioned stimuli CSs) when they promote the enactment of seeking habits in individuals with a SUD have not been elucidated. The present study aimed behaviorally and electrophysiologically to characterize the nature of the interaction between the reflexive and reflective systems recruited by CSs in individuals with a smoking habit. For this, we measured the behavioral performance and associated ERPs of 20 individuals with a smoking habit and 20 controls, who never smoked regularly, in a modified Go/NoGo task during which smoking-related CSs, appetitive, and neutral pictures, presented either in first-person or as a third-person visual perspective were displayed 250 ms before the Go/NoGo cue. We show that smoking-related cues selectively influence early incentive motivation-related attentional bias (N2 after picture onset), motor readiness and behavioral inhibition (Go-P3, NoGo-P3 and Pc) of individuals with a smoking habit only when presented from a first-person perspective. These data together identify the neural signature of the aberrant engagement of the reflexive and reflective systems during the recruitment of an incentive habit by CSs presented as if they had been response-produced, i.e., as conditioned reinforcers.

## Introduction

Substance use disorder (SUD) is a chronic relapsing disorder characterized by an aberrant motivation for, and the compulsive seeking and taking of, drugs (American Psychiatric Association, 2013). The transition in vulnerable individuals from recreational, controlled drug use to the persistent, relapsing engagement in drug seeking behavior despite adverse consequences that characterize addiction has been suggested to result from the development of maladaptive drug seeking habits (Belin & Everitt, 2010; Belin *et al*., 2013; Zilverstand *et al*., 2018; Fouyssac *et al*., 2022).

At the neural systems level, the progressive functional recruitment of the habit system over the course of the development of addiction has been shown to be reflected by a shift in the corticostriatal systems mediating the influence of drug-paired cues, or conditioned stimuli (CSs), on behavior from the ventral to the dorsolateral striatum (Vollstadt-Klein *et al*., 2010; Belin *et al*., 2013; Everitt & Robbins, 2016; Cox *et al*., 2017). Thus, while the motivational influence of Pavlovian CSs on behavior is associated with the activation of the Nucleus Accumbens (NAc) in recreational users, the same CSs functionally engage the dorsal striatum-dependent habit system in individuals with a long history of cocaine (Volkow *et al*., 2006; Cox *et al*., 2017), alcohol (Vollstadt-Klein *et al*., 2010), heroin (Xie *et al*., 2014) or nicotine use (McClernon *et al*., 2009; Detandt *et al*., 2017).

The functional engagement of the habit system in cue-provoked craving tasks has been shown to be the best predictor of actual long-term relapse (Zilverstand *et al*., 2018). This suggests that in individuals with a SUD, who suffer from an impaired *top-down control system* (Ramey & Regier, 2019; Luscher *et al*., 2020), drug-related stimuli recruit a bottom-up reflexive system (Bechara *et al*., 2006), which not only biases attention towards drug-paired CSs (attentional bias) (Field & Cox, 2008) but also facilitates foraging behavior (*Approach bias*) (Watson *et al*., 2012) and the engagement of instrumental drug seeking habits (Belin *et al*., 2013; Donamayor *et al*., 2021).

Attention bias (A.B.), behaviorally and electroencephalographically (EEG) characterized by faster response times (Wiers *et al*., 2013; Detandt *et al*., 2017) and by modulations of early (e.g., N1 and N2) and late (e.g., Late Positive Potential LPP, 400 – 600 ms) event-related components, respectively (Littel & Franken, 2007; Minnix *et al*., 2013; Rangaswamy & Porjesz, 2014; Robinson *et al*., 2016), has been suggested to reflect an implicit component of craving (Tiffany & Wray, 2012), that better predicts relapse (Goldstein & Volkow, 2011; McKay, 1999) and treatment efficacy than the explicit measures of craving (McKay, 1999; Goldstein & Volkow, 2011; Tiffany *et al*., 2012).

The loss of *top-down* control over these implicit mechanisms that contributes to the stimulus-bound and compulsive nature of the pursuit of the drug in individuals with a SUD (Everitt & Robbins, 2005; Everitt *et al*., 2008; Belin *et al*., 2013) has been related to alterations of the reflective system involving the dorsolateral prefrontal (DLPFC) and the anterior cingulate (ACC) cortices (Zilverstand *et al*., 2018). The neurophysiological signature of the early inhibitory control over prepotent responses exerted by these prefrontal cortical regions has been revealed in experimental tasks such as the Go/NoGo, the Stop-signal or the Stroop task as two inhibitory-specific ERP components: the N2 and the P3 (Kok *et al*., 2004). The N2 is a negative-going wave, with a peak occurring at frontocentral regions approximatively 200-300ms after stimulus onset, which has been frequently associated with the conflict monitoring function of the ACC (van Veen & Carter, 2002; Mathalon *et al*., 2003; Pandey *et al*., 2012). The P3, which culminates at frontocentral or parietal regions around 300 and 500ms after stimulus onset, has been related to the active motor inhibition process *per se* (Waller et al., 2021).

Thus, individuals with a smoking habit tend to evoke N2 (Luijten *et al*., 2011; see Pandey *et al*., 2012 for similar results with patients with an alcohol use disorder) and P3 (Evans *et al*., 2009; Luijten *et al*., 2014) of smaller amplitude than controls, in particular in the NoGo condition. In modified Go/NoGo paradigms in which a drug-paired CS is either presented just before a Go/NoGo signal or is the Go/NoGo signal itself, individuals with an addiction to alcohol, opiates or cocaine show higher rates of false alarms (Detandt *et al*., 2017), and a decrease in N2-NoGo and P3-NoGo amplitudes (Rangaswamy & Porjesz, 2014; Blanco-Ramos *et al*., 2019; Campanella *et al*., 2020). A smaller P3-NoGo, which characterizes SUD better than the N2 across different classes of drugs (Cohen *et al*., 1997; Fallgatter *et al*., 1998; Kamarajan *et al*., 2005; Colrain *et al*., 2011; for review, see Rangaswamy & Porjesz, 2014) (Luijten *et al*., 2014), is suggested to be a valuable neuromarker of the inhibitory deficits that predict relapse (Luijten *et al*., 2014).

Together these observations suggest that the engagement of the reflexive system by drug-paired CS challenges and exacerbates the weakness of the reflective system in individuals with SUD, thereby promoting rigid stimulus-bound behaviors. However, whether it is also the case for nicotine addiction remains to be established since some studies have reported larger Go/NoGo-P3 following drug-related compared to unrelated stimuli (Detandt *et al*., 2017), while others have reported behavioral (Yin *et al*., 2016) and/or neurophysiological evidence (Evans *et al*., 2009; Luijten *et al*., 2011; Buzzell *et al*., 2014; Liu *et al*., 2019) of a general deficit of the reflective system independent of a drug-paired CS recruitment of the reflexive system, e.g., when a CS is presented before the Go/NoGo cue. Thus, the nature of the interactions between the reflexive and reflective systems in nicotine addiction seems different from those involved in other SUDs. In the present study, we, therefore, sought behaviorally and neurophysiologically to characterize the nature of the interaction between reflexive and reflective processes in response to drug-paired CSs in individuals with a smoking habit.

To this aim, ERPs were measured in 20 individuals with an engrained smoking habit (S-group) and 20 control individuals who had no history of regular smoking (N-S group) during a modified Go/NoGo task in which pictures of smoking-related, appetitive, and neutral stimuli were displayed 250ms before a Go/NoGo trigger cue.

Since motivation-related processes occur about 300ms after stimulus onset (Robinson *et al*., 2016), the motivational component of the response to stimulus presentation (i.e., the LPP) was expected to be detected within the Go/NoGo-N2/P3 time window (Agudelo-Orjuela *et al*., 2021). Appetitive and smoking-related stimuli were expected to evoke a more positive LPP than neutral pictures in both groups. However, the attention grasping properties of pictures of smoking-related stimuli, reflected as a decrease of the N1/N2 amplitude (Rangaswamy & Porjesz, 2014), was expected to be observed only in the S-group, which was also predicted to evoke a more positive LPP than the NS-group (Littel & Franken, 2007; Minnix *et al*., 2013; Robinson *et al*., 2016). In accordance with an interactive reflexive-reflective system model, we expected an exacerbation of the difference between the Go/NoGo-N2 and/or the Go/NoGo-P3 between the S- and N.S.-group by the presentation of smoking-related as compared to neutral (Detandt *et al*., 2017) and appetitive stimuli, the latter being included in order to discriminate the specific effect of drug-paired CSs from that of general appetitive arousal on ERPs (Versace *et al*., 2017).

One important, often overlooked, property of many drug-paired CSs is that they are also response-produced, thereby acting as conditioned reinforcers, which support drug seeking habits over prolonged periods of time and contribute to the development of incentive habits (Olausson *et al*., 2004; Belin & Everitt, 2010; Belin *et al*., 2013; Fouyssac *et al*., 2022). Drug-paired CSs are, therefore, often experienced from a first-person perspective by individuals with SUD actively engaged in their drug seeking habits, thereby fostering the integration of addiction-specific multisensorial representations (Yalachkov *et al*., 2012). Accordingly, we manipulated how stimuli were presented in each picture, either from a first-person or a third-person perspective, which differentially recruit automatic embodiment (Canizales *et al*., 2013) and sensorimotor activation (Canizales *et al*., 2013; Galang *et al*., 2020). Accordingly, we expected the reflective system of individuals with a smoking habit to be relatively more challenged by the activation of the reflexive system by drug-paired CSs presented as first-person visual perspective compared to third-person visual perspective, reflected at the neurophysiological level by an increase of the P3-NoGo amplitude (Detandt *et al*., 2017). In these individuals with a smoking habit, we also expected that the reliance on Stimulus-Response association should facilitate response processing, as reflected by a decrease of the P3-Go amplitude (Rose *et al*., 2001), when both the drug- and task-related cues converge towards a similar response (Watson *et al*., 2012; Wiers *et al*., 2013; Detandt *et al*., 2017).

Beyond cue reactivity, the stimulus-bound response tendency characteristic of the S-group should also be reflected by modulations of the response-locked potentials evoked in the Go condition, that is, when no inhibition is required (Donkers & van Boxtel, 2004). Yet, these response-locked potentials, especially those evoked in correct Go trials, have not been thoroughly studied in individuals with SUD (Luijten *et al*., 2014; Luijten *et al*., 2016). There are two main ERP components following Go responses (Somon *et al*., 2019), the Correct Response Negativity (CRN) and the correct Positivity (Pc), representing an early (peak 80 ms after the response) and late (peak 300ms after the response) ERP, respectively. Falling in the Error-Related Negativity (ERN) time window, the CRN has been related to the evaluation of the accuracy of the response produced compared to the expected response. In contrast, the later Pc, considered a P300-like component related to the monitoring of the performance based on the representation of the outcome (i.e., subjective emotional significance, see Falkenstein *et al*., 2000). Accordingly, since the Pc, but not the CRN, is reduced when two stimuli call for a similar motor-response (Mathalon *et al*., 2002) we predicted that the Pc should be reduced selectively in the S-group when they show response facilitation brought about by the presentation of smoking-related CS in the Go condition, revealing the behavioral and neurophysiological signature of a drug-related habit.

## Materials and Methods

### Participants

Twenty individuals with a smoking habit (ISH) and twenty individuals who never regularly smoked (NRS, 15 women in each group) voluntary participated in this study, having signed an informed consent before the session. All participants were Spanish from Tenerife (Spain), and were recruited by convenience in the towns of La Laguna and Santa Cruz de Tenerife. They received 15€ after the completion of the study, which was approved by the Ethical Committee of the University of La Laguna (Tenerife, Spain).

Each participant first filled out an online questionnaire in which demographic and personal data were collected (gender, age, smoking history) and neuropsychological tests (Barratt Impulsiveness Scale (Salvo G & Castro S, 2013); STAI-R (Sandin *et al*., 1996); Beck’s depression questionnaire (Sanz *et al*., 2005); and the Positive Affect and Negative Affect Scale – PANAS (Watson *et al*., 1988) were self-administrated online using the Psytoolkit online software (Stoet, 2010). Smokers were also assessed for their smoking history (i.e., number of years of active smoking and number of cigarettes smoked per day) and their score in both the Spanish adaptation of the Fageström test (Becona & Vazquez, 1998) and the Obsessive-Compulsive Smoking Scale (Hitsman *et al*., 2010) in order to establish the severity of their smoking habit (see **Table 2**).

**Table 1.** Outcome of the evaluation (mean ± S.D.) of the emotional valence and arousal of pictures as a function of their type (appetitive, smoking-related or neutral) and the visual perspective in which they were taken (from a first- or third-person perspective).

| Type of pictures | Visual Perspective | Valence<br>(from 1 to 7) | Arousal<br>(from 1 to 7) |
| --- | --- | --- | --- |
| Appetitive | First-person | 5.4 ± 1.2 | 4.3 ± 1.5 |
|  | Third-Person | 5.5 ± 1.2 | 4.3 ± 1.5 |
| Smoking-related | First-person | 2.6 ± 1.4 | 4.1 ± 1.8 |
|  | Third-Person | 2.5 ± 1.3 | 4.0 ± 1.8 |
| Neutral | First-person | 4.4 ± 1.1 | 3.5 ± 1.3 |
|  | Third-Person | 4.6 ± 1.3 | 3.5 ± 1.3 |

**Table 2.**
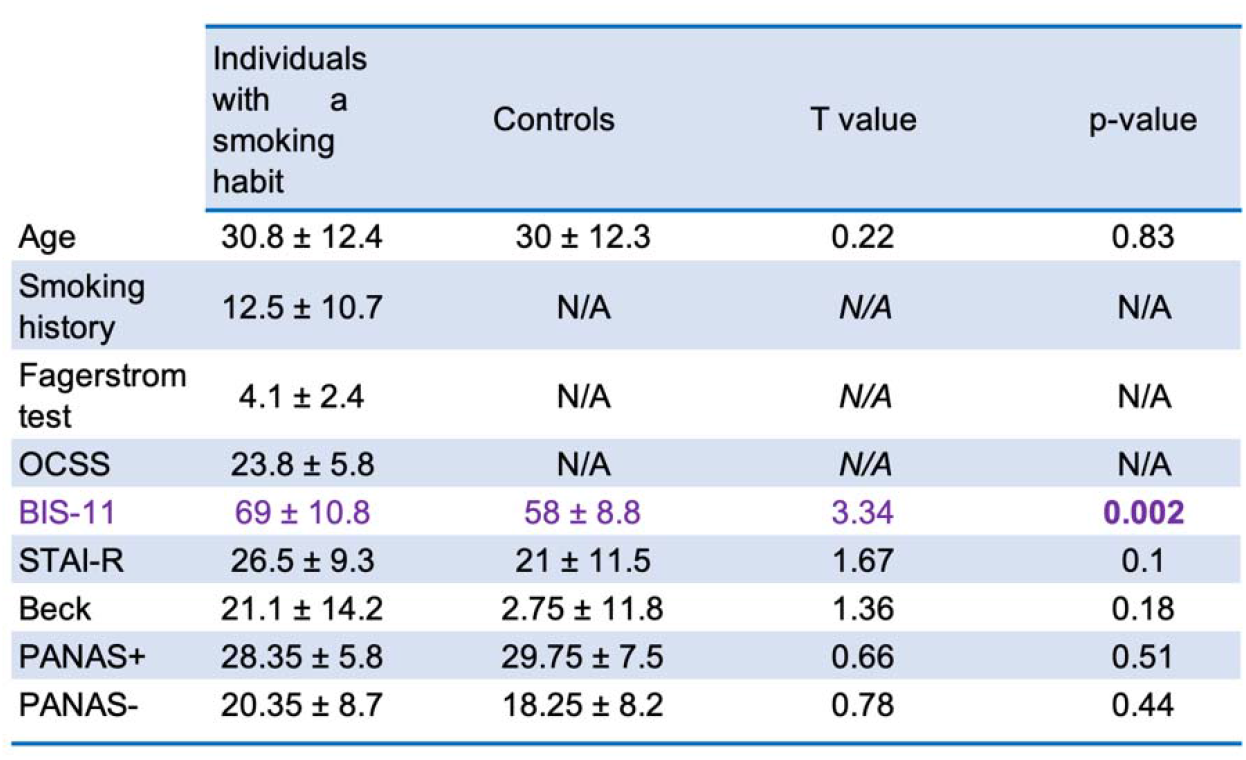
Demographic and personal data of individuals with a smoking habit and controls. (*Smoking history*; *Fagerström test*; *OCSS* = Obsessive Compulsive Smoking Scale; *BIS-11* = Barratt Impulsiveness Scale 11-items; *STAI-R* = State Trait Anxiety Inventory; *Beck* = Beck’s depression questionnaire; *PANAS+* and *PANAS-* = Positive Affect and Negative Affect Scale).

|  | Individuals with a smoking habit | Controls | T value | p-value |
| --- | --- | --- | --- | --- |
| Age | 30.8 ± 12.4 | 30 ± 12.3 | 0.22 | 0.83 |
| Smoking history | 12.5 ± 10.7 | N/A | N/A | N/A |
| Fagerstrom test | 4.1 ± 2.4 | N/A | N/A | N/A |
| OCSS | 23.8 ± 5.8 | N/A | N/A | N/A |
| BIS-11 | 69 ± 10.8 | 58 ± 8.8 | 3.34 | 0.002 |
| STAI-R | 26.5 ± 9.3 | 21 ± 11.5 | 1.67 | 0.1 |
| Beck | 21.1 ± 14.2 | 2.75 ± 11.8 | 1.36 | 0.18 |
| PANAS+ | 28.35 ± 5.8 | 29.75 ± 7.5 | 0.66 | 0.51 |
| PANAS- | 20.35 ± 8.7 | 18.25 ± 8.2 | 0.78 | 0.44 |

Before the beginning of the experimental session, each participant was then subjected to a measurement of their level of carbon monoxide (C.O.) using a Smokerlyzer (Bedfort Scientific Ltd., Rochester, UK). Then, each participant filled out the STAI-E and was given a short clinical interview led by J.D. in order to record any current medication and to detect any potential psychiatric, including other substance use, disorders using the MULTICAGE CAD-4 questionnaire (Pérez *et al*., 2007). A participant (control or smoker) who scored strictly above 2 (out of 4) in at least one of the eight sub-scales of the MULTICAGE CAD-4 (alcohol, illegal drugs, pathological gambling, Internet, video games, compulsive spending, eating disorders and sex addiction) was automatically excluded from the study. Three participants were excluded prior to the study (one presented a demyelinating disease, another one was under Xeristar 60 mg antidepressant treatment, and another one due to a suspected alcohol use disorder). Finally, the spontaneous level of craving of the individuals with a smoking habit was measured before and after the experimental session on a 31 points scale.

### Apparatus

Stimuli were presented using the E-Prime 2.0 software (PST) on a 17-inch monitor screen at a 768 × 1024 pixels resolution and 100Hz controlled by a P.C. The EEG signal was collected using the EasyCap system (BrainVision) equipped with 27 Ag/AgCl electrodes arranged in the international 10-20 system and referenced to the left mastoid. Four additional electrodes were used in order to provide bipolar recordings of the horizontal and vertical electrooculogram, two located at the outer canthus of each eye and two at the infraorbital and supraorbital regions of the right eye. The electrical activity was recorded and amplified with a bandwidth of 0.01–100 Hz and sampled at 500 Hz with impedances kept below 5 kΩ (electro-oculogram <10 kΩ).

Gaze position was also measured using an EyeLink 1000 system (S.R. Research Ltd., Ontario, Canada) recording at a 1000 Hz sampling rate, and synchronized to the E-Prime software. During the session, the participant sat at a distance of 70 cm from the screen, with the head resting on a chinrest adjusted to a comfortable position. A 7-point calibration was performed just before the initiation of the experiment.

### Materials

The two hundred and forty pictures created for this experiment were taken with a Canon professional camera by a member of the research team. Each photo was then post-processed in order to homogenize size and luminance. Each photo depicted a person interacting with one of three objects placed on a desk. Objects were chosen in order to create 3 sets of 80 pictures: smoking-related pictures presenting smoking-related objects (e.g., burning cigarette, lighter, cigarette pack, etc.), appetitive pictures (e.g., muffin, cookie, chocolate, etc.), and neutral pictures displaying office items and stationery (e.g., pen, book, paper sheets, etc.). Half the 80 pictures of each set were taken in third person visual perspective (i.e., facing the actor), the other half being taken in first person visual perspective. Examples of each stimulus are presented in **Figure 1**. The emotional valence and arousal-inducing properties of each picture were assessed and validated online on an independent pool of 83 participants (76% females, 15% of smokers) using the Psytoolkit software (Stoet, 2010).

**Figure 1.**
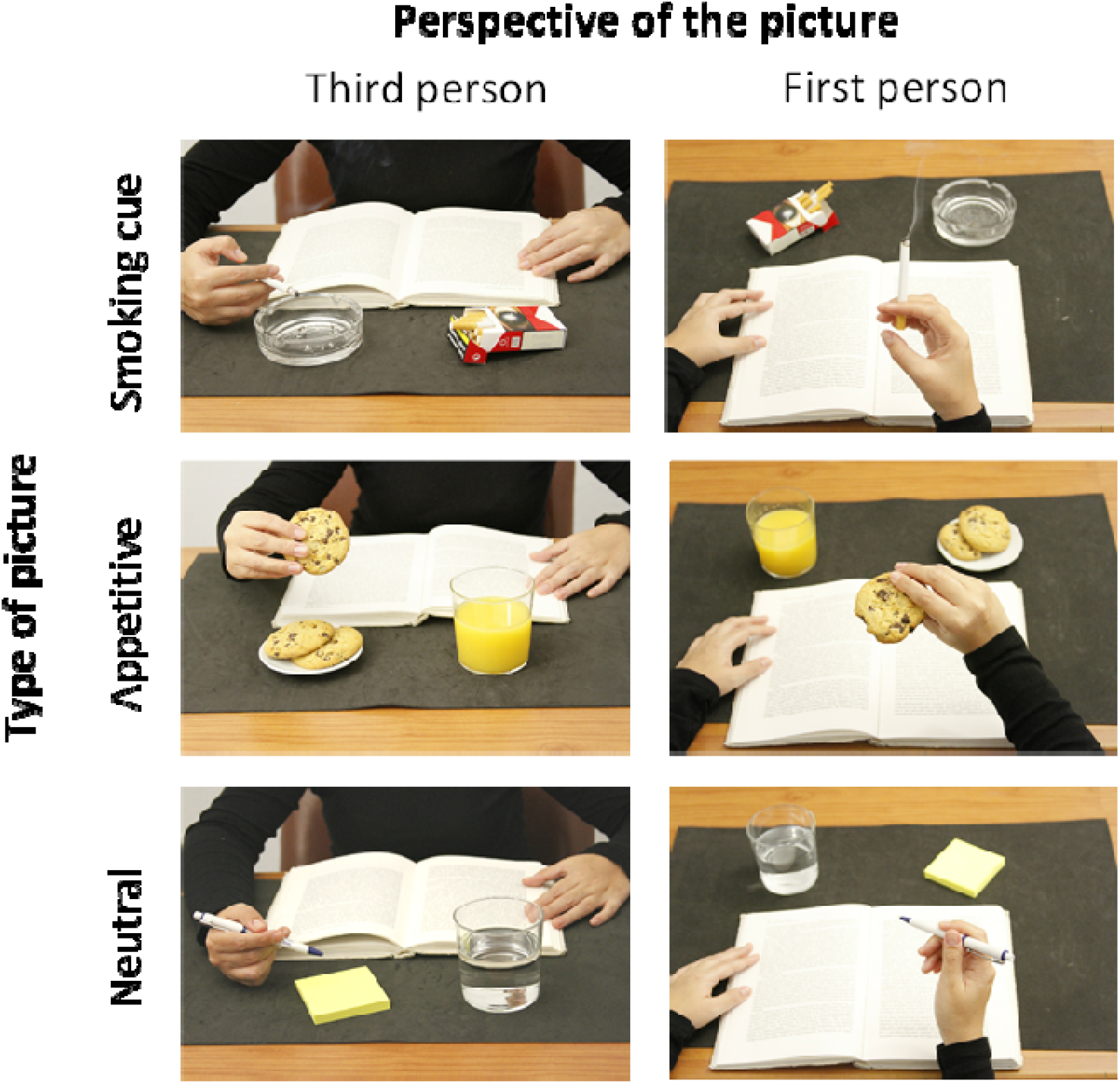
Example of smoking-related, appetitive or neutral pictures presented either from a first- or third-person visual perspective.

### Task paradigm and procedure

The G.O./NoGO task lasted approximately 30 minutes. A trial began with a grey screen for 450 ms, immediately followed by a 250 ms presentation of a picture (smoking-related, appetitive or neutral) displayed at the centre of the screen on a grey background. The frame of the picture was then coloured for 450 ms in blue or green (counterbalanced between subjects), indicating a Go or a NoGo trial after which the picture was replaced by a grey screen for 950 ms. Participants were instructed to respond in the Go trials by pressing the space bar of the keyboard with the right index as quickly and accurately as possible and to withhold that response in the NoGo trials. They were also told not to look at the border of the picture directly. Gaze position followed by an Eyelink 1000 (S.R. Research) was synchronized with the E-Prime software. If a participant’s gaze was detected to be directed at the frame of a picture the trial was immediately interrupted, and an error message appeared on screen. The cancelled trial was then presented again later in the experimental sequence.

There were 120 trials for each type of picture (80 in the Go condition and 40 in the NoGo condition) presented in random order in blocks (smoking-related, appetitive or neutral), the order of which was counterbalanced between participants. Participants were given the opportunity to rest for up to 5 minutes between each block.

### EEG preprocessing and data analysis

Data were processed offline using Brain Analyzer software (Brain Products GmbH, Gilching, Germany). The EEG recorded data were filtered offline with 0.1–30 Hz a band-pass, re-referenced to the algebraic mean of the activity at the two mastoids and corrected for ocular artefact using Independent Component Analysis (ICA, Makeig *et al*., 1995; Jung *et al*., 1998). Raw data, only including correct responses, were then segmented in 2-second epochs centred on the time of picture onset. A manual artefact rejection was performed, resulting in a similar trial rejection rate between individuals with a smoking habit and controls [3.5% and 3.2% respectively, *t*_*38*_ = 0.30, *p* = .77]. Then, in accordance with the objectives of the study, three re-segmentations were carried out:

1. on the picture onset in order to analyze the reflexive system-related N2 component (240 - 300 ms), which captures early motivation-related attentional mechanisms.
2. on the Go/NoGo cue onset in order to analyze the reflective system-related N2-Go/NoGo (250 - 300 ms) and the P3-Go/NoGo (310 - 370 and 420 - 470 ms).
3. on the (correct) responses in the Go condition in order to analyze response readiness and the associated post-response potential Pc (220 - 300 ms).

For each of these segmentations, the 200ms period preceding stimulus or response onset was used to calculate the baseline. Average ERP waves were computed for each segmentation, type of pictures and Go/NoGo cue.

The efficiency of the inhibitory mechanism in the Go/NoGo task was evaluated by the error rate and the response time (in milliseconds) of the participants. Response time was defined as the time elapsed between the appearance of the Go cue and the response (key press).

Repeated-measure Analyses of Variance (ANOVASs), implemented in “R” software (version 3.4.0) with the ULLRToolbox (https://sites.google.com/site/ullrtoolbox/home) were used, upon verification of assumption of normality and homogeneity of variance using the Shapiro-Wilk’s and Cochran test, respectively, to compare response times, the amplitude of the ERPs (ISH vs. NRS) as between-subject factor, and the type of cue (Go vs. NoGo), the type of picture (smoking-related, appetitive or neutral), the visual perspective (first vs. third perspective), the Regions (Frontal-left, Frontal-right, Frontocentral, Central-left, Central-right, Central, Parietal) and the Electrodes (e1, e2, e3) as within-subject factors.

Response times were first submitted to a log-transformation and then subjected to an ANOVA using the Group (ISH vs NRS) as between-participant factor, and the type of picture (smoking-related, appetitive or neutral) and the visual perspective (first vs third person point of view) as within-participant factors. Finally, false alarms and misses were analyzed by means of Chi-square tests (χ*2*).

The confirmation of significant main effects and differences among individual means were analyzed using the Newman-Keuls post-hoc test. For all analyses, significance was set at α= 0.05 and effect sizes were reported as partial eta squared (_p_η^2^) for every significant effect.

## Results

### Validation of the image bank

As expected the three picture categories evaluated by an independent cohort of 83 individuals were considered to differ in their emotional valence and arousing properties [main effect of picture: *F*_2,164_ = 118, *p* < 0.0001, _p_η^2^ = .59 and *F*_2,164_ = 8.43, *p* < 0.0005, _p_η^2^ = .09, respectively] (**Table 1**), with the pictures of appetitive stimuli deemed more appetitive [*t*_410_ = 7.79, *p* < 0.0001 and *t*_410_ = 23.05, *p* < 0.0001, respectively] and more arousing than those of neutral and smoking-related stimuli [*t*_410_ = 6.20, *p* < 0.0001 and *t*_410_ = 2.36, *p* = 0.02, respectively]. In contrast, pictures of smoking-related stimuli were deemed more arousing [*t*_410_ = 3.84, *p* = 0.003] but less appetitive than pictures depicting neutral stimuli [*t*_410_ = 15.26, *p* < 0.0001]. Even though visual perspective did not influence overall scoring of the image properties [*F*_1,82_ = 2.89, *p* = 0.09, *F*_1,82_ = 1.74, *p* = 0.19 for emotional valence and arousal property, respectively], it did have an effect, albeit small, on the judgement of the emotional valence [*F*_2,164_ = 4.68, *p* < 0.01, _p_η^2^ = .05], but not that of the arousing property [*F*_2,164_ = 1.66, *p* = 0.19], which was associated with a higher emotional valence score given to neural pictures presented as third-person [*t*_410_ = 2.61, *p* < 0.05, other *Ps* < .18] (**Table 1**).

### Characteristics of the experimental groups

Individuals with a smoking habit (ISH), who had a much higher level of carbon monoxide than control individuals (NRS) (14.05 ± 9.32 ppm vs 4.75 ± 1.45 ppm) [*F*_*1,38*_ = 18.76, *p* < 0.0001, _p_η^2^ = .33], were moderate smokers as determined by their scores in the Fagerström test and the OCSS (Wilson *et al*., 1999; Schane *et al*., 2010) (**Table 2**). ISH showed a characteristic increase in their spontaneous level of craving following the experimental session [16.2 ± 10.9 vs 18.42 ± 10.3, *F*_*1,19*_ = 6.23, *p* = 0.02, _p_η^2^ = .25] and, as expected (Bickel *et al*., 1999; Mitchell, 1999; Potvin *et al*., 2015), they were more impulsive than NRS, as revealed in their score in the BIS (p < .002). However, ISH did not differ from NRS in their levels of anxiety, mood or depression as assessed by the STAI, PANAS Beck, respectively (**Table 2**).

### Behavioral performance

ISH did not differ from NRS in their response time [*F*_1,38_ < 1, *p* = 0.87] which was not influenced by the type [*F*_2,37_ = 1.3, *p* = 0.27] or the visual perspective of the pictures [*F*_1,38_ = 2.8, *p* = 0.10]. Error rate (percentage of false alarms and omissions across trials) was very low and did not differ between ISH (3.9% and 0.7 respectively) and NRS (4.5% and 1.2 respectively) [χ*2* (1, *N* = 40) = 2.14, *p* = .14].

### ERPs reveal an electrophysiological signature of smoking habits

#### Picture onset

The amplitude of the picture onset-related P100 (80 – 110 ms) did not differ between ISH and NRS [main effect of group: *F*_1,38_ = 0.10, *p* = 0.75, group x visual perspective interaction: *F*_1,38_ = 3.51, *p* = 0.07, picture type x group interaction: *F*_1,38_ = 2.36, *p* = 0.11, picture type x visual perspective x group interaction: *F*_2,37_ = 0.51, *p* = 0.95]. However, pictures presented as first-person visual perspective evoked a more positive P100 than pictures as third-person visual perspective [*F*_1,38_ = 177.60, *p* < 0.0001, _p_η^2^ = .82] irrespective of the type of the picture [main effect of picture type: *F*_2,37_ = 0.69, *p* = 0.51, and visual perspective x picture type interaction: *F*_2,37_ = 0.24, *p* = 0.79] (**Figure 2A-D**). In contrast, the effect of the visual perspective of the picture on the amplitude of the N2 (240 – 300 ms) was determined by the electrode [*F*_2,37_ = 11.68, *p* = 0.0001, _p_η^2^ = .39], being greater in E1 than E2 and E3 [Follow-up ANOVA in First VP: *F*_2,76_ = 145.81, *p* < 0.0001, _p_η^2^ = .84, post-hoc comparisons: E1 vs E2: *t*_2356_ = 22.05, *p* ≤ 0.0001, E1 vs E3: *t*_2356_= 24.67, *p* = 0.0001, E2 vs E3: *t*_2356_= 2.62, *p* = 0.01 ; Follow-up ANOVA in Third VP: *F*_2,76_ = 128.62, *p* < 0.0001, _p_η^2^ = .87, post-hoc comparisons: E1 vs E2: *t*_2356_ = 23.62, *p* ≤ 0.0001, E1 vs E3: *t*_2356_= 25.32, *p* = 0.0001, E2 vs E3: *t*_2356_= 1.69, *p* = 0.10] and dependent on the group of participants [main effect of group: *F*_1,38_ = 0.67, *p* = 0.42, group x visual perspective interaction: *F*_1,38_ = 3.20, *p* = 0.08, and group x electrode x visual perspective interaction: *F*_2,76_ = 3.25, *p* = 0.04, _p_η^2^ = .13] as well as the valence of the picture [main effect of picture type: *F*_2,76_ = 1.74, *p* = 0.18, picture type x visual perspective interaction: *F*_2,76_ = 4.06, *p* = 0.02, _p_η^2^ = .19, and picture type x electrode x visual perspective interaction: *F*_4,152_ = 3.48, *p* < 0.01, _p_η^2^ = .24], resulting in a four-way interaction [*F*_4,35_ = 3.13, *p* = 0.03, _p_η^2^ = .26]. Between-group post-hoc comparisons revealed that the N2 was less negative in the ISH than the NRS only when the picture was smoking-related as third-person visual perspective [E1: *t*_46_ = 1.98, *p* ≤ 0.05; E2: *t*_46_ = 1.39, *p* = 0.25; E3: *t*_46_ = 1.22, *p* = 0.23; all other *Ps* > .10] (**Figure 2E-H**). Additionally, within-group post-hoc comparisons revealed that smoking-related pictures evoked a less negative N2 at the central electrode than neutral and appetitive pictures in ISH only (**Table 3**).

**Figure 2.**
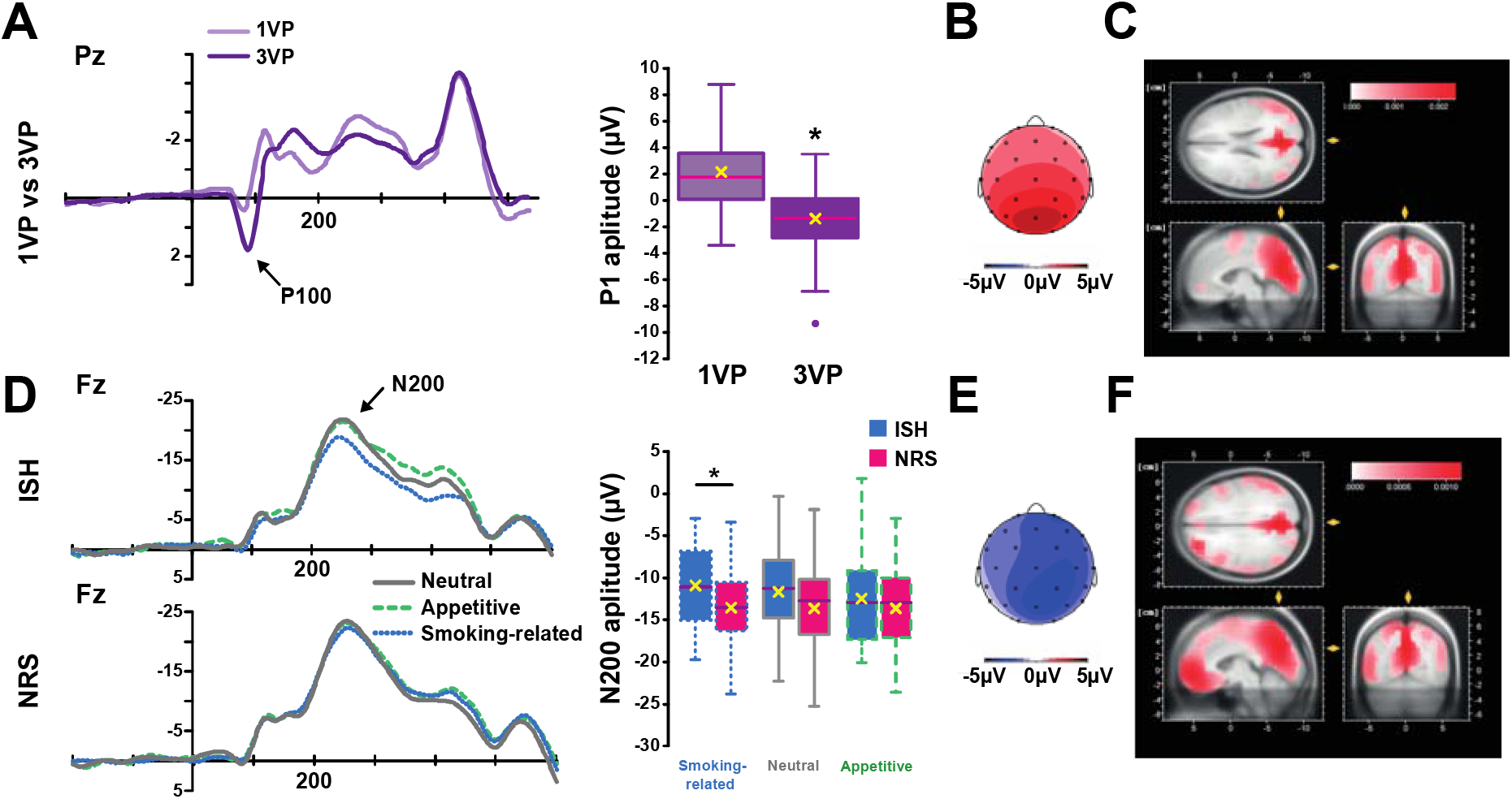
The N200 reflects selective attentional capture by drug-paired cues in ISH. **Top panel.** ERPs, and especially the P100, evoked in Pz by pictures from a first- (light purple) vs. third-person (dark purple) visual perspective (VP) (**A**) are represented alongside the P100 mean electrical voltage (µV) over the scalp (**B**) and its source estimation (LORETA) (**C**). The P100 evoked by pictures from a 3VP were more negative than those evoked by 1VP images. **Bottom panel**. ERPs, and especially the N200, evoked in Fz by neutral, appetitive and smoking-related pictures in ISH and NRS (**D**) are represented alongside the N200 mean electrical voltage (µV) in central electrodes (e1) (**E**) and its source estimation (LORETA) (**F**). ISH evoked a less negative N200 than NRS when presented with a smoking-related picture. No differences were observed between ISH and NRS in the N200 evoked by neutral and appetitive pictures. Note that source estimations of both P100 and N200 effects pinpoint to similar cortical areas (BA 31 and 7 respectively). *: p < 0.05

**Table 3.** Results of the post-hoc comparisons performed on the N2 amplitudes in smokers as a function of picture type (appetitive vs smoking-related vs. neutral), visual perspective (first-vs third-person), and electrode (E1 vs E2 vs E3). t and p values are reported.

| Electrode | Visual perspective | Comparison | T value | P value |
| --- | --- | --- | --- | --- |
| E1 | First-person | Appetitive - Smoking | -4.7 | < .0001 |
|  |  | Appetitive - Neutral | 0.22 | 0.83 |
|  |  | Smoking - Neutral | 4.92 | < .0001 |
|  | Third-person | Appetitive - Smoking | -3.77 | 0.0005 |
|  |  | Appetitive - Neutral | -3.54 | 0.0008 |
|  |  | Smoking - Neutral | 0.23 | 0.82 |
| E2 | First-person | Appetitive - Smoking | -2.8 | 0.01 |
|  |  | Appetitive - Neutral | 0.36 | 0.72 |
|  |  | Smoking - Neutral | 3.16 | 0.005 |
|  | Third-person | Appetitive - Smoking | -2.8 | 0.05 |
|  |  | Appetitive - Neutral | -2.32 | 0.05 |
|  |  | Smoking - Neutral | -0.04 | 0.97 |
| E3 | First-person | Appetitive - Smoking | -3.68 | < .0001 |
|  |  | Appetitive - Neutral | -0.05 | 0.96 |
|  |  | Smoking - Neutral | 3.63 | < .0001 |
|  | Third-person | Appetitive - Smoking | -2.84 | 0.01 |
|  |  | Appetitive - Neutral | -2.07 | 0.08 |
|  |  | Smoking - Neutral | 0.77 | 0.44 |

#### Go/NoGo cue onset

Analysis of the N2-Go/NoGo component (250 – 300 ms) (**Figure 3A**) confirmed that the NoGo cue evoked a more negative N2 than the Go cue [*F*_1,38_ = 62.99, *p* < 0.0001, _p_η^2^ = .62] while revealing that pictures in first-person visual perspective evoked a more negative N2 than pictures in third-person visual perspective [*F*_1,38_ = 4.88, *p* < 0.05, _p_η^2^ = .11]. These effects seemed to be independent of one another or of the picture type and experimental group as no other main effect, nor interaction were statistically significant.

**Figure 3.**
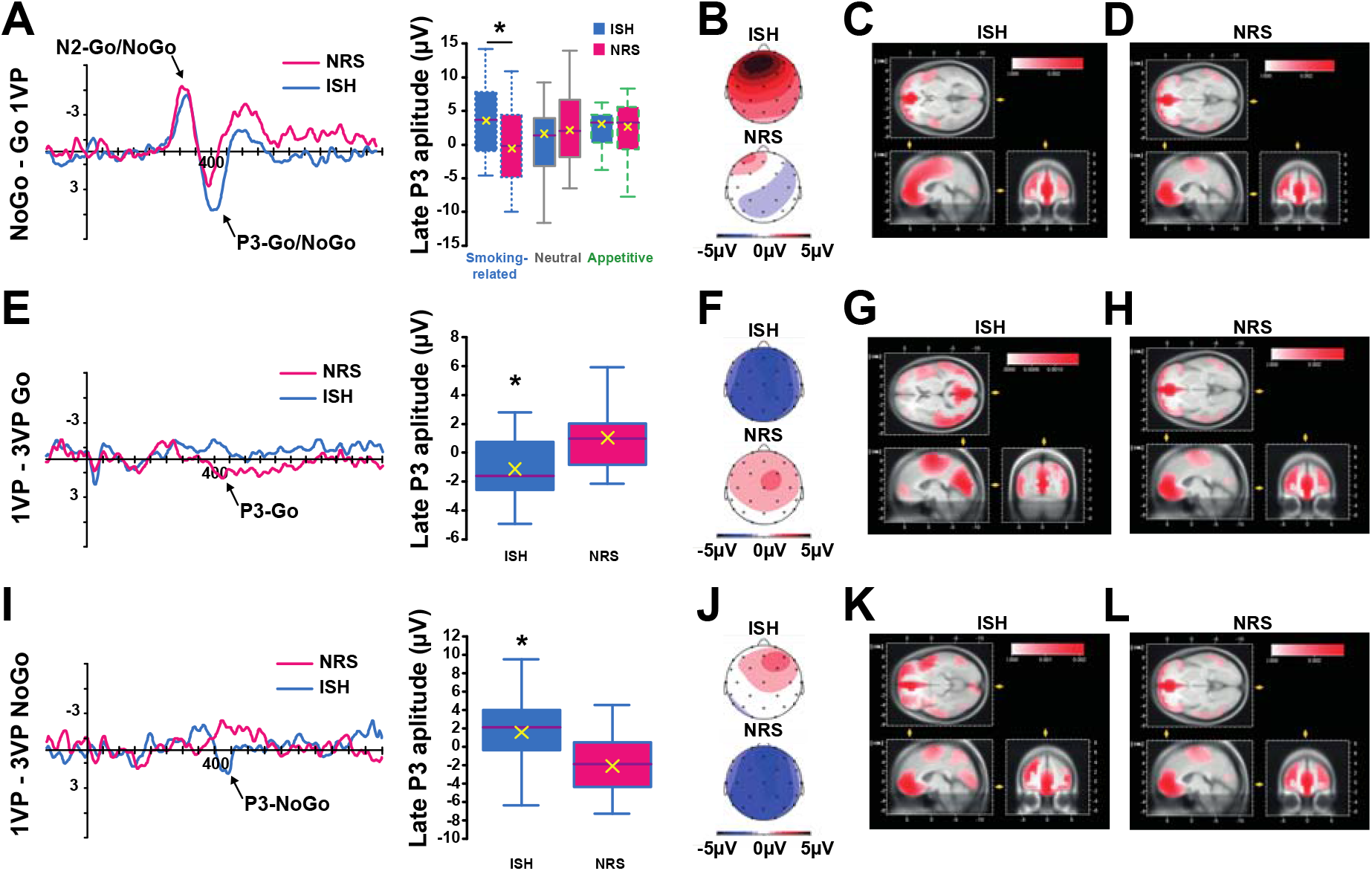
A late P3 selectively tracks the smoking-related cues presented from a first-person visual perspective in ISH. **(A)** The late P3 (420 – 470 ms) evoked at the F4 electrode following Go or NoGo cues differed between ISH and NRS when presented with smoking-related pictures from a first-person visual perspective (1VP). The electrical map (**B**) and source estimation in ISH (**C**) and NRS (**D**) revealed that the P3 was predominantly due to alterations in frontal modulations in both ISH and NRS (BA 10 and 32). (**E**) Subsequent analysis of the residual ERP evoked by smoking-related pictures in Go trials only revealed a differential impact of the visual perspective (the 1VP minus 3VP) in ISH as compared to NRS. The electrical map (**F**) and the source estimation in ISH (**G**) and NRS (**H**) revealed that this difference could be predominantly due to alterations in modulations of BA area 23 and are 10 in ISH and NRS, respectively. (**I**) In contrast, the residual ERP evoked by smoking-related pictures presented in NoGo trials revealed a reverse visual perspective effect (1VP minus 3VP) in each group compared to Go trials. The electrical map (**J**) and the source estimation in ISH (**K**) and NRS (**L**) revealed that the P3-NoGo presents a frontal distribution (BA 10 and 11) in both ISH and NRS. *: p< .05.

Analysis of the P3 component revealed a complex, group-dependent, time course of the onset of P3, which occurred later in ISH when presented with smoking-related pictures. Consequently, the P3 was analyzed over two different time windows, from 310 to 370 ms and from 420 to 470 ms after Go/NoGo cue onset.

The amplitude of the early P3 which was smaller when the picture was viewed as first- as compared to as third-person perspective [*F*_1,38_ = 6.09, *p* < 0.02, _p_η^2^ = .14], was influenced by the type of picture and the regions [picture type x region interaction: *F*_12,27_ = 2.90, *p* = 0.01, _p_η^2^ = .56, main effect of picture type: *F*_2,37_ = 1.73, *p* = 0.19, visual perspective x picture type: *F*_2,37_ = 0.16, *p* = 0.85, visual perspective x region: *F*_6,33_ = 5.33, *p* < 0.001, _p_η^2^ = .49, visual perspective x picture type x region interaction: *F*_12,27_ = 0.81, *p* = 0.64]. Post-hoc comparisons revealed that this was due to a smaller P3 evoked by appetitive pictures than neutral pictures in the frontocentral region only [*t*_9538_ = 3.39, *p* = 0.001] and a smaller P3 evoked by smoking-related pictures than neutral pictures in the frontocentral [*t*_9538_ = 4.80, *p* < 0.001], frontal right [*t*_9538_ = 2.48, *p* = 0.04], frontal left [*t*_9538_ = 3.5, *p* = 0.001] and central regions [*t*_9538_ = 3.36, *p* = 0.002]. The interaction between picture type and region was not dependent on having a smoking habit [main effect of group: *F*_1,38_ = 0.36, *p* = 0.55 and group x picture type x region: *F*_12,27_ = 1.37, *p* = 0.24].

The amplitude of the early P3 was also predicated on an interaction between the type of picture and Go/NoGo cue [*F*_2,37_ = 3.42, *p* = 0.04, _p_η^2^ = .15]. Post-hoc comparisons revealed that smoking-related and appetitive pictures evoked less positive P3 amplitudes than neutral pictures after the NoGo [*t*_9538_ = 9.03, *p* < 0.001 and *t*_9538_ = 5.66, *p* < 0.001, respectively] but not after the Go cue (all *p* > .10). In addition, the P3-NoGo evoked by the smoking-related pictures were less positive than that evoked by appetitive pictures [*t*_9538_ = 3.37, *p* = 0.001]. This effect was not a characteristic of a smoking habit since it was independent of the group of participants [main effect of group: *F*_1,38_ = 0.36, *p* = 0.55, and group x picture type x cue interaction: *F*_2,37_ = 1.15, *p* = 0.33].

The analysis of the late P3, in contrast, revealed a smoking-habit-specific electrophysiological signature. First, the late P3 amplitude was shown to be more positive in the NoGo than the Go condition [*F*_1,38_ = 4.43, *p* = 0.04, _p_η^2^ = .10], an effect that varied across regions [main effect of region: *F*_6,33_ = 11.78, *p* < 0.0001, _p_η^2^ = .68, and cue x region interaction: *F*_6,33_ = 5.42, *p* < 0.001, _p_η^2^ = .50], and as a function of the picture type [main effect of picture type: *F*_2,37_ = 0.38, *p* = 0.69, and cue x region x picture type interaction: *F*_12,27_ = 2.66, *p* = 0.02, _p_η^2^ = .54]. Thus, the Go/NoGo effect was stronger for smoking-related pictures than neutral pictures in the frontocentral region [*t*_4750_ = 2.33, *p* = 0.06] whereas it was stronger for appetitive pictures than for neutral pictures in the frontal right [*t*_4750_ = 2.58, *p* = 0.03], central [*t*_4750_ = 3.33, *p* = 0.02] and parietal regions [_*t*4750_ = 2.80, *p* = 0.01].

Finally, the Go/NoGo effect on the late P3 was dependent on whether individuals had a smoking habit and its interaction with the type of picture the visual perspective [group x cue x picture type x visual perspective interaction: *F*_2,37_ = 4.63, *p* = 0.01, _p_η^2^ = .20] (**Figure 3**). Post-hoc comparisons revealed that the Go/NoGo effect was stronger for ISH than for NRS only when the picture was smoking-related from a first-person visual perspective [*t*_42_= 2.80, *p* = 0.008, all other *p*s> 20] (**Figure 3A**).

In order to further characterize this effect, the amplitude of the late P3 evoked during Go and NoGo trials was analyzed separately, which revealed that the Go/NoGo cue effect was due an influence of visual perspective on the late P3-NoGo [*F*_1,38_ = 6.04, *p* = 0.02, _p_η^2^ = .14] and the late P3-Go [*F*_1,38_ = 7.05, *p* = 0.01, _p_η ^2^= .16] which were evoked differentially by ISH and NRS following presentation of smoking-related pictures. Post-hoc comparisons confirmed that smoking-related pictures when presented from a first-person compared to third-person visual perspective evoked a smaller P3-Go [*t*1558 = 6.56, *p* < .0001] in ISH but a more positive late P3-Go [*t*_1558_ = 2.38, *p* = .02] in NRS (**Figure 3E-H**). Conversely in NoGo trials, ISH produced a marginally more positive P3-NoGo [*t*_1558_ = 1.82, *p* = .07] with first compared to third-person visual perspective, while NRS who produced a less positive late P3-NoGo [*t*_1558_ = 9.33, *p* < .0001] in the presence of smoking-related pictures presented as first-person visual perspective as compared to when as third-person visual perspective (**Figure 3I-L**).

#### Response-locked potentials

Having established an ERP-based signature of smoking habits on inhibitory mechanisms, we investigated the RRPs in ISH and NRS over the 220 – 300 ms window that followed correct responses in the Go condition. The Pc amplitude was influenced by whether individuals had a smoking habit, the type of picture and its visual perspective [group x picture type interaction: *F*_2,37_ = 3.46, *p* = 0.04, _p_η^2^ = .16, and group x picture type x visual perspective interaction: *F*_2,37_ = 4.19, *p* = 0.02, _p_η^2^ = .18]. Post-hoc comparisons revealed that this was due to a less negative Pc evoked by ISH following a Go response for smoking-related pictures as compared to neutral [*t*_2356_ = 10.76, *p* < 0.0001] or appetitive pictures [*t*_2356_ = 12.29, *p* < 0.0001, all other *ps* > .13] (**Figure 4**). In addition, the visual perspective of the picture only influenced this effect in ISH [*F*_2,38_ = 3.61, *p* = 0.04, _p_η^2^ = .16] but not in NRS [*F*_2,38_ = 1.53, *p* = 0.23], thereby revealing an important role of the visual perspective in the integration the motivational value of the response in ISH. Thus, in ISH responses for smoking-related images presented from a first-person visual perspective evoked a larger Pc than for both neutral [*t*_2356_ = 5.36, *p* < 0.0001] and appetitive pictures [*t*_2356_ = 5.61, *p* < 0.0001], which did not differ from each other [*t*_2356_ = 0.25, *p* = 0.81]. This effect was attributed to modulation of the frontal cortex (Brodmann area, BA 6) by source estimation (**Figure 4A-D**). In contrast, responses made for smoking-related pictures presented from a third-person visual perspective evoked a more positive Pc than for smoking-related pictures presented from a first-person visual perspective [*t*_2356_ = 2.63, *p* = 0.01]. This larger smoking picture related response-evoked Pc is also greater than that evoked for neutral [*t*_2356_ = 14.43, *p* < 0.0001] and appetitive pictures [*t*_2356_ = 11.11, *p* < 0.0001]. However, responses made for appetitive images presented from a first-person visual perspective evoked a larger Pc than those made for neutral pictures [*t*_2356_ = 3.32, *p* < 0.001] (**Figure 4E-H**). Source estimation indicated that these effects were all related to modulations of the orbitofrontal cortex (BA10).

**Figure 4.**
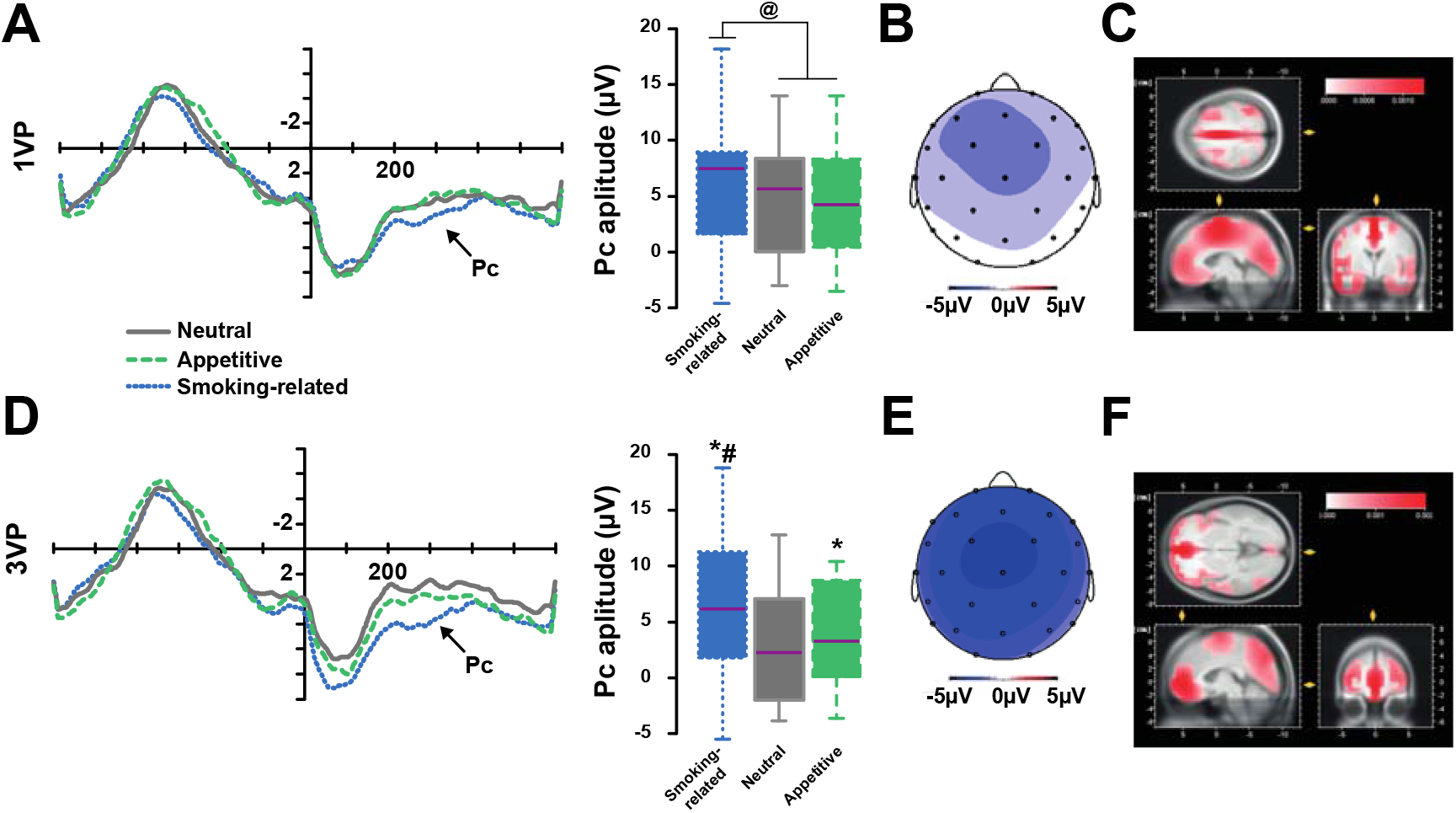
The motor response-locked Pc in Go trial is greater in ISH for smoking-related pictures than appetitive and neutral pictures. (**A-C**) The amplitude of the Pc (in Cz) evoked by ISH tracked only the difference between responses made with with smoking-related pictures presented from a first-person visual perspective (1VP) and those made with appetitive and neutral pictures. The latter were associated with a response that evoked a similar Pc with a lower amplitude than that evoked by the former. Source estimation indicated that these effects were related to modulations of the frontal lobe (BA6) (**B&C**). (**D**) The Pc evoked by ISH following a response made with a smoking related picture from a 3VP, which was of a greater amplitude that that evoked by the same pictures from a 1VP, was also greater than that evoked by responses made with appetite and neutral pictures. However, when presented from a 3VP appetitive pictures were associated with responses evoking greater Pc than neutral pictures. Source estimation indicated that these effects were related to modulations of the orbitofrontal cortex (BA10) (**E&F**). @: p < 0.05; *: different from neural, p < 0.05; #: different from appetitive, p < 0.05.

## Discussion

ISH performed as well as NRS overall in the modified Go/NoGo task developed for this study, both groups making very few errors and showing fast response times, both well within the standard range reported for this type of task (Evans *et al*., 2009; Buzzell *et al*., 2014). Thus, ISH involved in the present study, while having a smoking habit, did not display global impairment of their inhibitory system in line with the outcome of previous studies (Evans *et al*., 2009; Buzzell *et al*., 2014) at odds with evidence of poorer inhibition, i.e, lower accuracy in No-Go trials only, in ISH with Fagerström score similar to that of those in the present study (Luijten *et al*., 2011). This apparent discrepancy may stem from differences in the experimental design, such as the absence of the non-smoking-related appetitive stimuli or pictures presented from a first-person visual perspective that were used in the present study, or from a relative volatility of behavioral measures such as accuracy to capture inhibitory control deficits.

Performance in the task was associated with classical N2 and P3 ERP components, the magnitude of which was predicated on the nature of the trial (Go vs NoGo cue) as previously described (Kok *et al*., 2004; Randall & Smith, 2011), but not influenced by having a smoking habit. Drug-cue processing has also been related to a modulation of late ERP components, in particular the LPP, a positive component usually recorded in centroparietal regions between 300 and 600 ms after stimulus onset (Littel & Franken, 2007; Minnix *et al*., 2013; Robinson *et al*., 2016). The differential modulation of the LPP by drug-related cues as compared to drug-unrelated cues (Littel & Franken, 2007; Minnix *et al*., 2013; Robinson *et al*., 2016) has long been suggested to represent differential cognitive processing of the emotional or motivational signification of the stimulus. For example, Versace et al. (2011) reported more LPP of greater positive amplitudes when evoked by smoking-related, pleasant and unpleasant stimuli than those evoked by neutral stimuli. We expected to detect the motivational component of the response to cue presentation in the range of the LPP, that is, in the P3 (310 – 370 ms) taking the Go/NoGo cue as stimulus onset (Agudelo-Orjuela *et al*., 2021). As anticipated, smoking-related and appetitive pictures evoked a less positive P3 (310 – 370 ms, that is 560 ms after the onset of the pictures) than neutral pictures in both ISH and NRS, thereby demonstrating that the two groups processed the emotional characteristics of the pictures similarly.

However, in individuals with a SUD, drug-related cues generate specific multisensory representations associated with overt and covert motivational mechanisms, including approach bias (Watson *et al*., 2012). Importantly, many smoking-related cues are response-produced, and therefore perceived from a first-person visual perspective (e.g., automatic embodiment, see Canizales *et al*., 2013; Galang *et al*., 2020), by ISH, in whom they act as conditioned reinforcers. Conditioned reinforcers not only bridge delays to reinforcement, but they also facilitate the development of incentive habits and the subsequent development of compulsive drug seeking (Belin & Everitt, 2010; Belin *et al*., 2013; Fouyssac *et al*., 2022). The specific influence of conditioned reinforcers on motivational processes has been illustrated by the greater influence cues exert on behavior when presented as first-person visual perspective than third person visual perspective (Yalachkov *et al*., 2012). Accordingly, we manipulated the visual perspective of each stimulus, which influenced in first instance the amplitude of the P100 in both ISH and NRS. Similar perspective effects have been reported (Rigato *et al*., 2019) and consistently related to somatosensory processing involved in representations of oneself and others. In addition to the general ability of the visual perspective to influence early ERP components, smoking-related pictures evoked a N2 of smaller amplitude than neutral and appetitive pictures, only when presented from a first-person visual perspective. This observation is in agreement with the long-established modulation of early ERP components (e.g., N1 and N2; for a review, see Rangaswamy & Porjesz, 2014) by drug-related cues in individuals with a SUD which has been interpretated as a drug-specific attentional bias. Descriptive source estimation revealed in the present study that these effects originated respectively from the Posterior cingulate cortex (Brodmann area -BA-31) and the parietal superior cortex (BA7), areas that have been related to the processing of the spatial information in goal-oriented behaviors (Hadjidimitrakis *et al*., 2019) and associated CS-induced attentional bias (Engelmann *et al*., 2009).

Beyond the processing of the motivational value of cues, this study also identified the nature of the influence of these cues on inhibition processing. Thus, while the emotionally loaded appetitive and smoking-related pictures evoked a smaller early P3-NoGo (310 – 370) than neutral pictures in both ISH and NRS, a well-established signature the impact of the emotional characteristics of stimuli on inhibition (Agudelo-Orjuela *et al*., 2021), this was followed by a smaller late P3-Go (420 – 470 ms), in ISH only, evoked by smoking-related compared to neutral pictures, and only when presented from a first-person visual perspective. Smoking-related pictures presented from a first-person visual perspective may, therefore, provoke a transient impairment of inhibitory processes associated with an automatic recruitment of the incentive motivational processes related to the influence of conditioned reinforcers on the expression of incentive habits (Belin *et al*., 2013; Jones *et al*., 2018). However, follow-up analysis showed that the modulation of the late P3-Go/NoGo (420 – 470 ms) by smoking-related cues was not only due to a decrease of the P3-Go amplitude in ISH, but also to an increase of that of the P3-NoGo. ISH produced a smaller P3-Go in the presence of a smoking-related pictures from a first-as compared to third-person visual perspective, a profile opposite to that shown by NRS and suggested by source estimation to be associated with an activation of the posterior cingulate cortex (B.A. 23) the prefrontal cortex (BA10) in the former and the latter, respectively. This decrease in the P3-Go amplitude shown by ISH may represent the impact of previously activated incentive habit-related Stimulus-Response rules by smoking-related cues presented from a first-person visual perspective (i.e. attentional bias, N2 effect at picture onset), resulting in the facilitation of response processing when both the drug-related and task-related cues point or converge towards a similar response (Watson *et al*., 2012; Wiers *et al*., 2013; Detandt *et al*., 2017). Since NRS have never learnt about, or indeed experienced response-produced smoking-related cues, it would be expected that they did not show such facilitation of response processing by these stimuli when presented from a first-person visual perspective, which to some extent may even be alien to them. It was therefore not surprising that NRS displayed the exact opposite neurophysiological signature to that shown by ISH, e.g., NRS evoked a larger P3-Go than ISH when presented with a smoking-related picture as first-person visual perspective, suggesting that such unfamiliar situation requires more response processing than that associated with appetitive images.

The specific inhibition processing profile of smoking habits was associated with a unique response potential signature. The analysis of the response potentials in Go trials revealed that ISH evoked a smaller, less negative response potential than NRS for responses associated with a smoking-related picture as compared to neutral or appetitive pictures irrespective of the visual perspective in which they are presented. The time course of this effect (220 and 300 ms) is in line with the P300-like correct Positivity component (Pc). Even though the nature of the cognitive function it reflects remains relatively elusive (Bates *et al*., 2004), the Pc is increasingly considered to reflect a post-response monitoring process sensitive to the subjective value of the outcome (Falkenstein *et al*., 2000) and to the congruency between responses in that it tends to decrease when two stimuli call for a similar response (Mathalon *et al*., 2002). Accordingly, the decrease in the Pc amplitude observed in the present study in ISH when responding following the presentation of a smoking-related cue suggests a facilitation of the post-response evaluation processing reflective of the recruitment of ingrained automatic, outcome value independent, responding mediated by stimulus-response associations (Robbins & Costa, 2017) associated here with activation of Sensory-Motor-Area (SMA, BA6) revealed by source estimation when the response is given after the presentation of a smoking-related picture from a first-person visual perspective.

While the association of the posterior cingulate cortex with the smaller P3-Go evoked by ISH in response to a smoking-related picture from a first person visual perspective discussed above may point towards an influence of the perception of any stimulus presented as first-person visual perspective as being more related to the self and thereby influencing the level of familiarity or affective salience (Murray *et al*., 2012) of the image (Zilverstand *et al*., 2018), the source estimation in the SMA of the smaller Pc evoked ISH responding following the presentation of smoking-related pictures as first-person visual perspective suggests a specific recruitment of automatic responding mediated by an action knowledge network (Yalachkov *et al*., 2009; Yalachkov & Naumer, 2011). In addition, while only responses made by ISH following a smoking-related picture evoked a lower Pc than neutral pictures when presented as first-person visual perspective, responding following both appetitive and smoking-related pictures evoked a smaller Pc than following neutral pictures when presented from a third-person visual perspective, reflective of a facilitation of post-response evaluation that is proportional to the emotional value of the stimulus when presented from a third-person visual perspective. In line with this interpretation, the differential Pcs evoked for responses following presentations of stimuli from a third-person visual perspective did not originate in the SMA, as shown for Pc evoked by responses made following smoking-related pictures presented as first-person visual perspective, but instead in the orbitofrontal cortex (BA10), which has been related to decision-making processes based on stimulus–reward mapping (Young & Shapiro, 2011).

## Conclusion

The influence of drug-, and in particular alcohol-, related CSs on inhibitory processing in individuals with a SUD has been well documented (for a meta-analysis, see Jones *et al*., 2018), however that of smoking-related cues, especially when presented in the same visual perspective as when response produced, in individuals with a smoking habit remained to be fully characterized. By comparing the influence of smoking-related cues presented either as first- or third-person visual perspective to that of drug-unrelated appetitive pictures on the neurophysiological correlates of inhibitory processing (Versace *et al*., 2017), and response monitoring (Falkenstein *et al*., 2000; Mathalon *et al*., 2002) this study revealed a selective signature of the engagement of the inhibitory system and automatic response processing by smoking related pictures in ISH.

Together the results of this experiment suggest that in ISH, who have a long history of drug foraging under the control of the conditioned reinforcing properties of response-produced smoking-related cues, thence experienced as first-person visual perspective, smoking-related cues presented in the same visual perspective as a conditioned reinforcer recruit a relatively weak inhibitory process while facilitating the recruitment of habitual responding.

## Authors’ contribution

J.D. and D.B. designed the experiments. J.D. conducted the experiments and the analysis of the data. J.D. and D.B. wrote the manuscript.

## Acknowledgements

This work was conducted at Laboratory of Cognitive Neuroscience and Psycholinguistics, University of La Laguna. J.D. was supported by a Postdoctoral fellowship from the University of La Laguna (Programa Agustin de Betancourt). HAB was supported by the Spanish Ministry of Science and Innovation (grant PID2020-118487GB-I00). D.B. was supported by a Medical Research Council Programme grant (MR/N02530X/1) to DB, Barry Everitt, Amy Milton, Jeffrey Dalley and Trevor Robbins.

For the purpose of open access, the author has applied a Creative Commons Attribution (CC BY) licence to any Author Accepted Manuscript version arising.

The data can be available at: https://docs.google.com/spreadsheets/d/1VyYDZckcA6-z_CRX7YnK98MfoW_OX7Dd/edit?usp=sharing&ouid=113770177571978399708&rtpof=true&sd=true

## References

Agudelo-Orjuela, P., de Vega, M. & Beltran, D. (2021) Mutual influence between emotional language and inhibitory control processes. Evidence from an event-related potential study. Psychophysiology, 58, e13743.

Association, A.P. (2013) The Diagnostic and Statistical Manual of Mental Disorders: DSM 5. bookpointUS.

Bates, A.T., Liddle, P.F., Kiehl, K.A. & Ngan, E.T.C. (2004) State dependent changes in error monitoring in schizophrenia. Journal of Psychiatric Research, 38, 347–356.

Bechara, A., Noel, X. & Crone, E.A. (2006) Loss of willpower: abnormal neural mechanisms of impulse control and decision making in addiction. In Wiers, R.W., Stacy, A.W. (eds) Handbook of implicit cognition and addiction. Sage, Thousand Oaks, CA, pp. 215–232.

Becona, E. & Vazquez, F.L. (1998) The Fagerstrom Test for Nicotine Dependence in a Spanish sample. Psychol Rep, 83, 1455–1458.

Belin, D., Belin-Rauscent, A., Murray, J.E. & Everitt, B.J. (2013) Addiction: failure of control over maladaptive incentive habits. Curr Opin Neurobiol, 23, 564–572.

Belin, D. & Everitt, B.J. (2010) The Neural and Psychological Basis of a Compulsive Incentive Habit. In Steiner, H., tseng, K. (eds) Handbook of basal ganglia structure and function. Elsevier, ACADEMIC PRESS, pp. 571–592.

Bickel, W.K., Odum, A.L. & Madden, G.J. (1999) Impulsivity and cigarette smoking: delay discounting in current, never, and ex-smokers. Psychopharmacology, 146, 447–454.

Blanco-Ramos, J., Cadaveira, F., Folgueira-Ares, R., Corral, M. & Rodriguez Holguin, S. (2019) Electrophysiological Correlates of an Alcohol-Cued Go/NoGo Task: A Dual-Process Approach to Binge Drinking in University Students. Int J Environ Res Public Health, 16.

Buzzell, G.A., Fedota, J.R., Roberts, D.M. & McDonald, C.G. (2014) The N2 ERP component as an index of impaired cognitive control in smokers. Neurosci Lett, 563, 61–65.

Campanella, S., Schroder, E., Kajosch, H., Hanak, C., Veeser, J., Amiot, M., Besse-Hammer, T., Hayef, N. & Kornreich, C. (2020) Neurophysiological markers of cue reactivity and inhibition subtend a three-month period of complete alcohol abstinence. Clin Neurophysiol, 131, 555–565.

Canizales, D.L., Voisin, J.I., Michon, P.E., Roy, M.A. & Jackson, P.L. (2013) The influence of visual perspective on the somatosensory steady-state response during pain observation. Front Hum Neurosci, 7, 849.

Cohen, H.L., Porjesz, B., Begleiter, H. & Wang, W. (1997) Neurophysiological correlates of response production and inhibition in alcoholics. Alcohol Clin Exp Res, 21, 1398–1406.

Colrain, I.M., Sullivan, E.V., Ford, J.M., Mathalon, D.H., McPherson, S.L., Roach, B.J., Crowley, K.E. & Pfefferbaum, A. (2011) Frontally mediated inhibitory processing and white matter microstructure: age and alcoholism effects. Psychopharmacology (Berl), 213, 669–679.

Cox, S.M.L., Yau, Y., Larcher, K., Durand, F., Kolivakis, T., Delaney, J.S., Dagher, A., Benkelfat, C. & Leyton, M. (2017) Cocaine Cue-Induced Dopamine Release in Recreational Cocaine Users. Sci Rep, 7, 46665.

Detandt, S., Bazan, A., Quertemont, E. & Verbanck, P. (2017) Smoking addiction: the shift from head to hands: Approach bias towards smoking-related cues in low-dependent versus dependent smokers. J Psychopharmacol, 31, 819–829.

Donamayor, N., Ebrahimi, C., Garbusow, M., Wedemeyer, F., Schlagenhauf, F. & Heinz, A. (2021) Instrumental and Pavlovian Mechanisms in Alcohol Use Disorder. Current Addiction Reports, 8, 156–180.

Donkers, F.C. & van Boxtel, G.J. (2004) The N2 in go/no-go tasks reflects conflict monitoring not response inhibition. Brain Cogn, 56, 165–176.

Engelmann, J.B., Damaraju, E., Padmala, S. & Pessoa, L. (2009) Combined effects of attention and motivation on visual task performance: transient and sustained motivational effects. Front Hum Neurosci, 3, 4.

Evans, D.E., Park, J.Y., Maxfield, N. & Drobes, D.J. (2009) Neurocognitive variation in smoking behavior and withdrawal: genetic and affective moderators. Genes Brain Behav, 8, 86–96.

Everitt, B.J., Belin, D., Economidou, D., Pelloux, Y., Dalley, J.W. & Robbins, T.W. (2008) Review. Neural mechanisms underlying the vulnerability to develop compulsive drug-seeking habits and addiction. Philos Trans R Soc Lond B Biol Sci, 363, 3125–3135.

Everitt, B.J. & Robbins, T.W. (2005) Neural systems of reinforcement for drug addiction: from actions to habits to compulsion. Nat Neurosci, 8, 1481–1489.

Everitt, B.J. & Robbins, T.W. (2016) Drug Addiction: Updating Actions to Habits to Compulsions Ten Years On. Annu Rev Psychol, 67, 23–50.

Falkenstein, M., Hoormann, J., Christ, S. & Hohnsbein, J. (2000) ERP components on reaction errors and their functional significance: a tutorial. Biol Psychol, 51, 87–107.

Fallgatter, A.J., Wiesbeck, G.A., Weijers, H.G., Boening, J. & Strik, W.K. (1998) Event-related correlates of response suppression as indicators of novelty seeking in alcoholics. Alcohol Alcohol, 33, 475–481.

Field, M. & Cox, W.M. (2008) Attentional bias in addictive behaviors: a review of its development, causes, and consequences. Drug Alcohol Depend, 97, 1–20.

Fouyssac, M., Pena-Oliver, Y., Puaud, M., Lim, N.T.Y., Giuliano, C., Everitt, B.J. & Belin, D. (2022) Negative Urgency Exacerbates Relapse to Cocaine Seeking After Abstinence. Biol Psychiatry, 91, 1051–1060.

Galang, C.M., Jenkins, M. & Obhi, S.S. (2020) Exploring the effects of visual perspective on the ERP components of empathy for pain. Soc Neurosci, 15, 186–198.

Goldstein, R.Z. & Volkow, N.D. (2011) Dysfunction of the prefrontal cortex in addiction: neuroimaging findings and clinical implications. Nat Rev Neurosci, 12, 652–669.

Hadjidimitrakis, K., Bakola, S., Wong, Y.T. & Hagan, M.A. (2019) Mixed Spatial and Movement Representations in the Primate Posterior Parietal Cortex. Front Neural Circuits, 13, 15.

Hitsman, B., Shen, B.J., Cohen, R.A., Morissette, S.B., Drobes, D.J., Spring, B., Schneider, K., Evans, D.E., Gulliver, S.B., Kamholz, B.W., Price, L.H. & Niaura, R. (2010) Measuring smoking-related preoccupation and compulsive drive: evaluation of the obsessive compulsive smoking scale. Psychopharmacology (Berl), 211, 377–387.

Jones, A., Robinson, E., Duckworth, J., Kersbergen, I., Clarke, N. & Field, M. (2018) The effects of exposure to appetitive cues on inhibitory control: A meta-analytic investigation. Appetite, 128, 271–282.

Jung, T.-P., Makeig, S., Bell, A.J. & Sejnowski, T.J. (1998) Independent Component Analysis of Electroencephalographic and Event-Related Potential Data. In Poon, P.W.F., Brugge, J.F. (eds) Central Auditory Processing and Neural Modeling. Springer US, Boston, MA, pp. 189–197.

Kamarajan, C., Porjesz, B., Jones, K.A., Choi, K., Chorlian, D.B., Padmanabhapillai, A., Rangaswamy, M., Stimus, A.T. & Begleiter, H. (2005) Alcoholism is a disinhibitory disorder: neurophysiological evidence from a Go/No-Go task. Biol Psychol, 69, 353–373.

Kok, A., Ramautar, J.R., De Ruiter, M.B., Band, G.P. & Ridderinkhof, K.R. (2004) ERP components associated with successful and unsuccessful stopping in a stop-signal task. Psychophysiology, 41, 9–20.

Littel, M. & Franken, I.H.A. (2007) The effects of prolonged abstinence on the processing of smoking cues: an ERP study among smokers, ex-smokers and never-smokers. Journal of Psychopharmacology, 21, 873–882.

Liu, C., Dong, F., Li, Y., Ren, Y., Xie, D., Wang, X., Xue, T., Zhang, M., Ren, G., von Deneen, K.M., Yuan, K. & Yu, D. (2019) 12 h Abstinence-Induced ERP Changes in Young Smokers: Electrophysiological Evidence From a Go/NoGo Study. Front Psychol, 10, 1814.

Luijten, M., Kleinjan, M. & Franken, I.H. (2016) Event-related potentials reflecting smoking cue reactivity and cognitive control as predictors of smoking relapse and resumption. Psychopharmacology (Berl), 233, 2857–2868.

Luijten, M., Littel, M. & Franken, I.H.A. (2011) Deficits in Inhibitory Control in Smokers During a Go/NoGo Task: An Investigation Using Event-Related Brain Potentials. Plos One, 6.

Luijten, M., Machielsen, M.W., Veltman, D.J., Hester, R., de Haan, L. & Franken, I.H. (2014) Systematic review of ERP and fMRI studies investigating inhibitory control and error processing in people with substance dependence and behavioural addictions. J Psychiatry Neurosci, 39, 149–169.

Luscher, C., Robbins, T.W. & Everitt, B.J. (2020) The transition to compulsion in addiction. Nat Rev Neurosci, 21, 247–263.

Makeig, S., Bell, A.J., Jung, T.-P. & Sejnowski, T.J. (1995) Independent component analysis of electroencephalographic data Proceedings of the 8th International Conference on Neural Information Processing Systems. MIT Press, Denver, Colorado, pp. 145–151.

Mathalon, D.H., Fedor, M., Faustman, W.O., Gray, M., Askari, N. & Ford, J.M. (2002) Response-monitoring dysfunction in schizophrenia: an event-related brain potential study. J Abnorm Psychol, 111, 22–41.

Mathalon, D.H., Whitfield, S.L. & Ford, J.M. (2003) Anatomy of an error: ERP and fMRI. Biol Psychol, 64, 119–141.

McClernon, F.J., Kozink, R.V., Lutz, A.M. & Rose, J.E. (2009) 24-h smoking abstinence potentiates fMRI-BOLD activation to smoking cues in cerebral cortex and dorsal striatum. Psychopharmacology (Berl), 204, 25–35.

McKay, J.R. (1999) Studies of factors in relapse to alcohol, drug and nicotine use: a critical review of methodologies and findings. J Stud Alcohol, 60, 566–576.

Minnix, J.A., Versace, F., Robinson, J.D., Lam, C.Y., Engelmann, J.M., Cui, Y., Brown, V.L. & Cinciripini, P.M. (2013) The late positive potential (LPP) in response to varying types of emotional and cigarette stimuli in smokers: a content comparison. Int J Psychophysiol, 89, 18–25.

Mitchell, S.H. (1999) Measures of impulsivity in cigarette smokers and non-smokers. Psychopharmacology, 146, 455–464.

Murray, R.J., Schaer, M. & Debbane, M. (2012) Degrees of separation: a quantitative neuroimaging meta-analysis investigating self-specificity and shared neural activation between self- and other-reflection. Neurosci Biobehav Rev, 36, 1043–1059.

Olausson, P., Jentsch, J.D. & Taylor, J.R. (2004) Repeated nicotine exposure enhances responding with conditioned reinforcement. Psychopharmacology (Berl), 173, 98–104.

Pandey, A.K., Kamarajan, C., Tang, Y., Chorlian, D.B., Roopesh, B.N., Manz, N., Stimus, A., Rangaswamy, M. & Porjesz, B. (2012) Neurocognitive deficits in male alcoholics: An ERP/sLORETA analysis of the N2 component in an equal probability Go/NoGo task. Biological Psychology, 89, 170–182.

Pérez, P.E., Monje, M., Alonso, F., Giron, M., Lopez, M. & Romero, J. (2007) Validacion de un instrumento para la deteccion de trastornos de control de impulsosy adicciones: el MULTICAGE CAD-4 Trastornos Adictivos, pp. 269–279.

Potvin, S., Tikasz, A., Dinh-Williams, L.L., Bourque, J. & Mendrek, A. (2015) Cigarette Cravings, Impulsivity, and the Brain. Front Psychiatry, 6, 125.

Ramey, T. & Regier, P.S. (2019) Cognitive impairment in substance use disorders. CNS Spectr, 24, 102–113.

Randall, W.M. & Smith, J.L. (2011) Conflict and inhibition in the cued-Go/NoGo task. Clin Neurophysiol, 122, 2400–2407.

Rangaswamy, M. & Porjesz, B. (2014) Understanding alcohol use disorders with neuroelectrophysiology. Handb Clin Neurol, 125, 383–414.

Rigato, S., Bremner, A.J., Gillmeister, H. & Banissy, M.J. (2019) Interpersonal representations of touch in somatosensory cortex are modulated by perspective. Biol Psychol, 146, 107719.

Robbins, T.W. & Costa, R.M. (2017) Habits. Curr Biol, 27, R1200–R1206.

Robinson, M.J., Fischer, A.M., Ahuja, A., Lesser, E.N. & Maniates, H. (2016) Roles of “Wanting” and “Liking” in Motivating Behavior: Gambling, Food, and Drug Addictions. Curr Top Behav Neurosci, 27, 105–136.

Rose, M., Verleger, R. & Wascher, E. (2001) ERP correlates of associative learning. Psychophysiology, 38, 440–450.

Salvo G, L. & Castro S A. (2013) Confiabilidad y validez de la escala de impulsividad de Barratt (BIS-11) en adolescentes. Revista chilena de neuro-psiquiatría, 51, 245–254.

Sandin, B., Chorot, P. & McNally, R.J. (1996) Validation of the spanish version of the Anxiety Sensitivity Index in a clinical sample. Behav Res Ther, 34, 283–290.

Sanz, J., García Vera, M.P., Espinosa, R., Fortún, M. & Vázquez Valverde, C. (2005) Adaptación española del Inventario para la Depresión de Beck-II (BDI-II): 3. Propiedades psicométricas en pacientes con trastornos psicológicos. Clínica y Salud, 16, 121–142.

Schane, R.E., Ling, P.M. & Glantz, S.A. (2010) Health effects of light and intermittent smoking: a review. Circulation, 121, 1518–1522.

Somon, B., Campagne, A., Delorme, A. & Berberian, B. (2019) Evaluation of performance monitoring ERPs through difficulty manipulation in a response-feedback paradigm. Brain Research, 1704, 196–206.

Stoet, G. (2010) PsyToolkit: a software package for programming psychological experiments using Linux. Behav Res Methods, 42, 1096–1104.

Tiffany, S.T., Friedman, L., Greenfield, S.F., Hasin, D.S. & Jackson, R. (2012) Beyond drug use: a systematic consideration of other outcomes in evaluations of treatments for substance use disorders. Addiction, 107, 709–718.

Tiffany, S.T. & Wray, J.M. (2012) The clinical significance of drug craving. Ann N Y Acad Sci, 1248, 1–17.

van Veen, V. & Carter, CS (2002) The anterior cingulate as a conflict monitor: fMRI and ERP studies. Physiol Behav, 77, 477–482.

Versace, F., Engelmann, J.M., Deweese, M.M., Robinson, J.D., Green, C.E., Lam, C.Y., Minnix, J.A., Karam-Hage, M.A., Wetter, D.W., Schembre, S.M. & Cinciripini, P.M. (2017) Beyond Cue Reactivity: Non-Drug-Related Motivationally Relevant Stimuli Are Necessary to Understand Reactivity to Drug-Related Cues. Nicotine Tob Res, 19, 663–669.

Versace, F., Minnix, J.A., Robinson, J.D., Lam, C.Y., Brown, V.L. & Cinciripini, P.M. (2011) Brain reactivity to emotional, neutral and cigarette-related stimuli in smokers. Addict Biol, 16, 296–307.

Volkow, N.D., Wang, G.J., Telang, F., Fowler, J.S., Logan, J., Childress, A.R., Jayne, M., Ma, Y. & Wong, C. (2006) Cocaine cues and dopamine in dorsal striatum: mechanism of craving in cocaine addiction. J Neurosci, 26, 6583–6588.

Vollstadt-Klein, S., Wichert, S., Rabinstein, J., Buhler, M., Klein, O., Ende, G., Hermann, D. & Mann, K. (2010) Initial, habitual and compulsive alcohol use is characterized by a shift of cue processing from ventral to dorsal striatum. Addiction, 105, 1741–1749.

Waller, D.A., Hazeltine, E. & Wessel, J.R. (2021) Common neural processes during action-stopping and infrequent stimulus detection: The frontocentral P3 as an index of generic motor inhibition. Int J Psychophysiol, 163, 11–21.

Watson, D., Clark, L.A. & Tellegen, A. (1988) Development and validation of brief measures of positive and negative affect: the PANAS scales. J Pers Soc Psychol, 54, 1063–1070.

Watson, P., de Wit, S., Hommel, B. & Wiers, R.W. (2012) Motivational Mechanisms and Outcome Expectancies Underlying the Approach Bias toward Addictive Substances. Front Psychol, 3, 440.

Wiers, C.E., Kuhn, S., Javadi, A.H., Korucuoglu, O., Wiers, R.W., Walter, H., Gallinat, J. & Bermpohl, F. (2013) Automatic approach bias towards smoking cues is present in smokers but not in ex-smokers. Psychopharmacology (Berl), 229, 187–197.

Wilson, D., Parsons, J. & Wakefield, M. (1999) The health-related quality-of-life of never smokers, ex-smokers, and light, moderate, and heavy smokers. Prev Med, 29, 139–144.

Xie, C., Shao, Y., Ma, L., Zhai, T., Ye, E., Fu, L., Bi, G., Chen, G., Cohen, A., Li, W., Chen, G., Yang, Z. & Li, S.J. (2014) Imbalanced functional link between valuation networks in abstinent heroin-dependent subjects. Mol Psychiatry, 19, 10–12.

Yalachkov, Y., Kaiser, J., Gorres, A., Seehaus, A. & Naumer, M.J. (2012) Smoking experience modulates the cortical integration of vision and haptics. Neuroimage, 59, 547–555.

Yalachkov, Y., Kaiser, J. & Naumer, M.J. (2009) Brain regions related to tool use and action knowledge reflect nicotine dependence. J Neurosci, 29, 4922–4929.

Yalachkov, Y. & Naumer, M.J. (2011) Involvement of action-related brain regions in nicotine addiction. J Neurophysiol, 106, 1–3.

Yin, J., Yuan, K., Feng, D., Cheng, J., Li, Y., Cai, C., Bi, Y., Sha, S., Shen, X., Zhang, B., Xue, T., Qin, W., Yu, D., Lu, X. & Tian, J. (2016) Inhibition control impairments in adolescent smokers: electrophysiological evidence from a Go/NoGo study. Brain Imaging Behav, 10, 497–505.

Young, J.J. & Shapiro, M.L. (2011) Dynamic coding of goal-directed paths by orbital prefrontal cortex. J Neurosci, 31, 5989–6000.

Zilverstand, A., Huang, A.S., Alia-Klein, N. & Goldstein, R.Z. (2018) Neuroimaging Impaired Response Inhibition and Salience Attribution in Human Drug Addiction: A Systematic Review. Neuron, 98, 886–903.

